# Bacteriophage infection reveals pre-emptive cooperative dormancy in stationary phase *Escherichia coli*

**DOI:** 10.64898/2026.09.23.753752

**Authors:** Pavel A. Ivanov, Olga Yu Timoshina, Sergey V. Mureev, Maria A. Letarova, Andrey V. Letarov

## Abstract

Bacterial cultures entering stationary phase (SP) undergo complex physiological alterations, helping the cells to survive when medium resources are exhausted. The onset of various stress responses underlying SP physiology is generally believed to be due to reactions of individual cells to the environmental cues such as starvation, toxic metabolites, etc. The SP physiological state makes bacteria unsuitable for the replication of most bacteriophages; however, some phages are able to infect and multiply in them. We investigated *Escherichia coli* phage DH23 growth in SP cultures of *E. coli* MG1655 infected at different hours post inoculation (ages). Under our conditions the cells enter SP at about 6 h, but the phage replication was possible till 11h and then dropped abruptly by 13h of culture aging, indicating an abrupt physiological change to deeper dormancy during SP. The onset of this phage-inhibiting middle-SP dormancy (MSPD) turned out to be mediated by intercellular communication mediated by the middle-SP signal particles (MSP) which are larger than 100 kDa and contain both RNA and DNA. These MSP are inactivated by RNAse or by DNAse, enabling phage growth in 14h-old cultures and lifts the tolerance of such cultures to kanamycin. Moreover, the RNAse or DNAse treatment makes 24h culture spent media suitable for additional cell growth, and cultures initiated in fresh LB supplemented with RNAse reach an OD600 about 1.5 times higher compared to the untreated control. This indicates that MSP signaling influences cell physiology well before the MSPD onset and helps the population cease growth pre-emptively to save about 1/3 of medium resources to support the viability during SP.

## Introduction

The growth of bacteria in batch culture is a self-limiting process, in which a short period of rapid multiplication, including what is commonly referred to as logarithmic growth phase, is followed by slowing down and complete arrest of culture growth when resources in the medium become depleted. Then, during a period lasting for a few days, generally termed stationary phase (SP), most bacteria do not divide but remain viable (reviewed in [1]). The vast majority of the SP cells are able to rapidly resume growth and division if diluted into fresh media. By contrast, a small fraction can show delayed multiplication (persister cells) or lose the ability to restart division altogether while retaining the membrane integrity. The latter are so-called viable but non-culturable cells (VBNC) [2–4]. Stationary phase is followed by dying off of the vast majority of the cells, with stabilization of surviving bacteria at a much lower level, which is described as the long-term stationary phase or post-stationary phase, and which last for months or even years [5–7].

*Escherichia coli* growing under optimal conditions (37°C with vigorous aeration) in rich LB media is one of the most widely used models for stationary phase physiology research [1,7–10]. Although many studies employ more defined synthetic media such as M9 [11,12], the LB model appears to be more relevant to the conditions faced *in vivo* by a population of pathogenic organisms during chronic infections in the human or animal organism, where nutrients are more diverse than in a single carbon source synthetic medium. A slowing down of the growth in LB medium is normally seen at about 4-5 h post dilution of the overnight culture and by 6-7 h the culture riches it’s maximum density entering SP. The transition to stationary phase in LB or in minimal media is not merely a growth arrest because of resource exhaustion, but a complex physiological stress response (termed general stress response) helping the bacterial population to survive a period of starvation. This response includes global transcription reprogramming orchestrated by the alternative sigma factor RpoS [13,14], and alterations of both translation and transcription because of the accumulation of the cellular alarmone (p)ppGpp, known as the stringent response [15,16]. The onset of the general stress response is also associated with sequestration of the majority of ribosomes in non-active dimers by the proteins RMF and HPF [17–19]. Finally, in deep SP (around 16-18h post inoculation) the formation of large protein aggregates at the cell poles is observed [20]. These aggregates sequester a large number of the proteins involved in energy metabolism, macromolecular syntheses, and cell growth and division machineries ([20], see also review [21]), thereby lowering the metabolic activity of the cell. The maturation and structural compaction of the polar aggregates have been demonstrated to correlate with the depth of cell dormancy (SP cell ⇒ persister ⇒ VBNC) [3].

Transition between log-phase and SP is also associated with changes in cell morphology. The balanced growth in which the cell elongation is tightly coordinated with division gives way to so-called reductive division, leading to shortening of the cells [1,22,23]. Later during SP, cells undergo so called dwarfing which further reduces the cell size due to the turnover of part of the cytoplasm and internal membrane material alongside the shedding of a fraction of the outer membrane (OM) in the form of the outer membrane vesicles (OMV) [1,24]. Both reductive division and dwarfing drive the elongated log-phase *E. coli* cells to become smaller and ovoid during SP.

All of the physiological changes at the entry into SP and during this period are controlled by multiple regulatory circuits mainly acting at the post-transcriptional level, involving not only regulatory proteins but also multiple small RNAs, adaptors and anti-adaptors controlling the degradation of RpoS and other key proteins and other mechanisms [3,17,18]. However, to our knowledge, all the SP physiology circuits described so far act inside the cell responding to the environmental cues sensed by each cell individually. Moreover, in another proteobacterium, *Xanthomonas oryzae*, quorum-sensing-coordinated social behaviors were reported to be suppressed upon the entry into SP [25].

The activation of SP cells upon addition of the nutrients is less understood compared to the general stress response, although significant progress has been made over the last decade (see [26] for review). Here again the response of the cells to more favorable conditions, e.g., the addition of nutrients, is, to our knowledge, deemed individual.

Most bacteriophages are not able to propagate on host cultures that have entered SP or even at the transition between log-phase and SP. Although the inability of the phages to grow in cultures that have attained a certain “critical density” was recognized more than 100 years ago [27], the exact mechanisms of this restriction are poorly understood. One of the most common manifestations of the phage growth restriction in SP (or pre-SP) cells is the phenomenon of the bacteriophage plaque expansion arrest when the bacterial lawn reaches its maximal density [28]. However, some phages were shown to be able to productively infect the SP cells [29–34] and even long-starving cells termed deeply dormant [11]. Among *E. coli* bacteriophages such a stationary phase infective (SPI) phenotype has long been known for phage T7. Recently we analyzed the phenomenology of T7 growth on mature bacterial lawns [35] and in SP liquid cultures of *E. coli* K-12 MG1655 [36] and confirmed that the unlimited plaque expansion phenotype is indeed linked to the infectivity against the SP cells. We also characterized a new SPI coliphage *Dhillonvirus* DH23 featuring a plaque growth phenotype very similar to that of T7 [37].

Here we investigated in depth the interactions of the phage DH23 with the cultures of *E. coli* MG1655 at different growth phases and demonstrated that the ability of the phage DH23 to multiply in such cultures fell down abruptly already during the SP between 11h and 13h of incubation, delimiting the early and middle SP. We demonstrated that the timely onset and maintenance of cell dormancy corresponding to middle SP requires an extracellular signal produced by the cells. This signal is a large particle exceeding 100 kDa proteins in size and containing both RNA and DNA. Removal or inactivation of the signal dramatically changes the outcome of the culture infections by DH23, T7 and other SPI phages. We also demonstrate that cooperative dormancy mediated by the extracellular signal particles takes place before the exhaustion of the medium resources (nutrients and capacity), saving about 40% of these resources for cell maintenance.

## Results

### DH23 infection outcomes in the cultures at different growth phases

Under our conditions the cultures reached maximal density by 6 h (Fig S1).

**Fig. S1.**
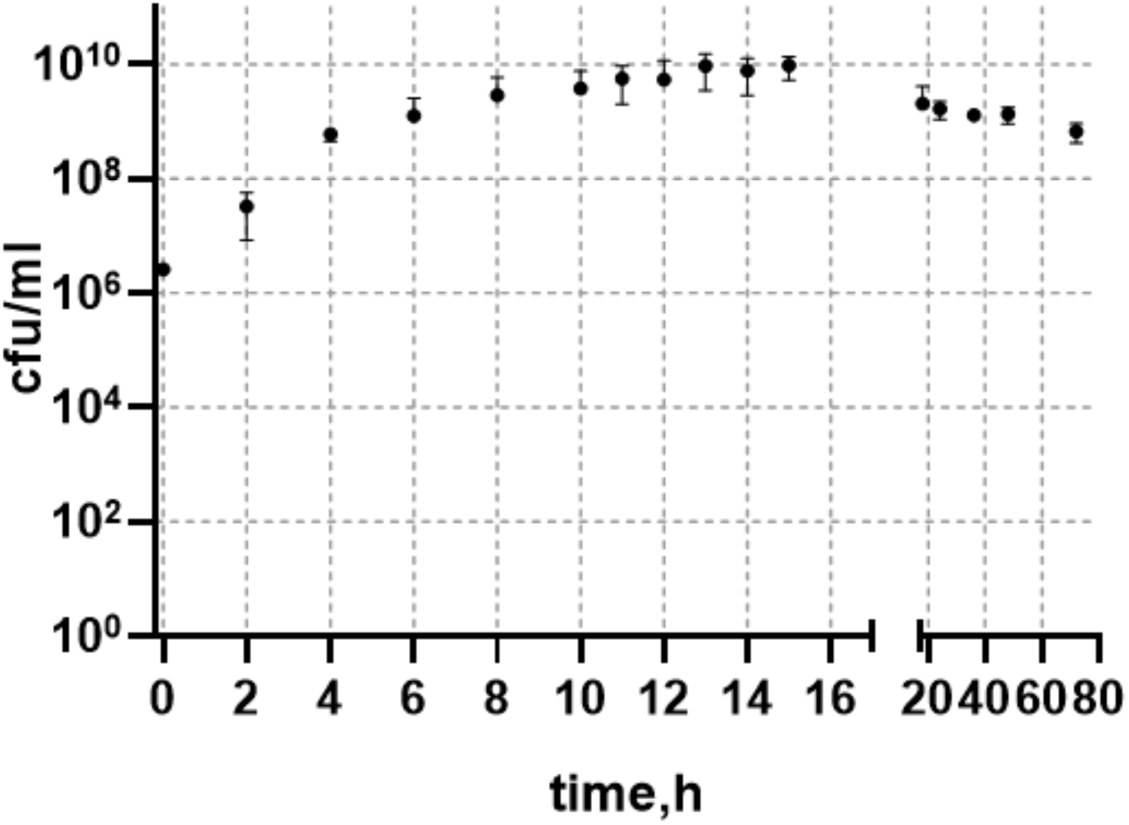
Growth curve of *E. coli* MG1655 strain.

The aliquots of the cultures from different time points were inoculated at a multiplicity of infection (MOI) of 0.1 and incubated for 24 h at 37°C with agitation. The resulting CFU counts at time points 2h and 4 h were identical to the non-infected control, then dropped ca. 2 orders of magnitude for the points 5h – 10h and rose again to the level of the control in the cultures older than 12 h (Fig.1). At the early time points (2h and 4h) clear lysis was observed within 3-4 h post infection, but then the re-growth of the resistant mutants led to a secondary rise in culture density up to the control level. The cells present in the infected 2h and 4h cultures were DH23-resistant as confirmed by the phage stock titration on the subclones (n=3). At the time points 5h – 11h most of the cells were lysed by the phage but the surviving sub-population was not able to re-grow, presumably because of the exhaustion of the medium resources by the time of infection. At later time points the phage failed to lyse the cultures or reduce the CFU counts. The total PFU counts (the sum of free phages and infected cells) remained high until 11 h, but dropped starting from 13 h of aging to the level of the initially added phage or even slightly lower in cultures older than 16h (Fig. 1).

**Fig. 1.**
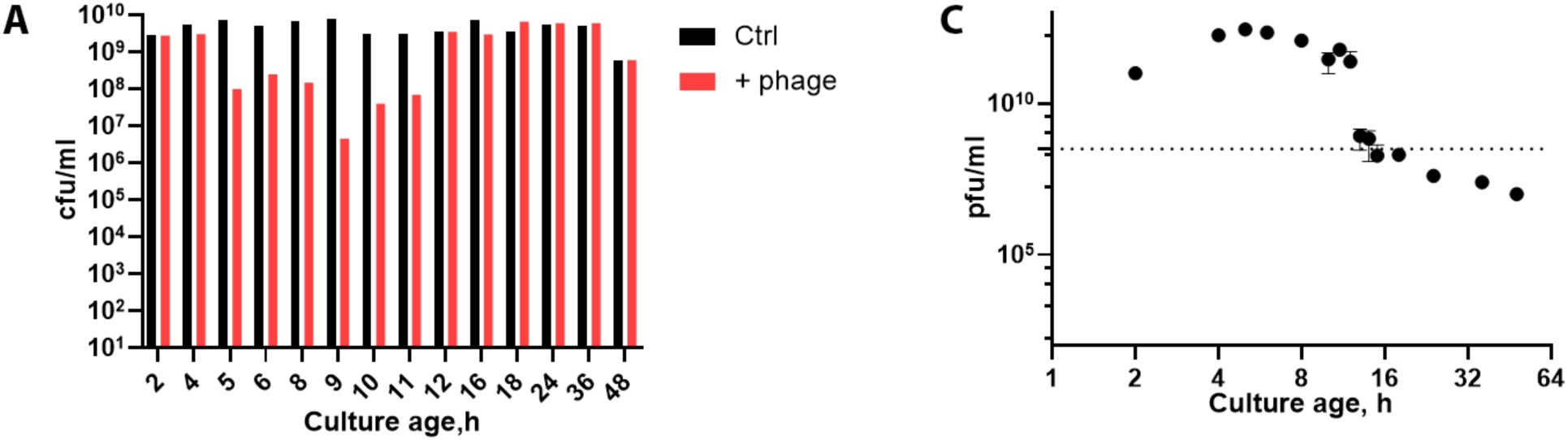
Infection of cultures aged 2 – 48h with DH23 phage. CFU counts (**A**), and phage titer (**B**) determined 24h post infection. The dotted line indicates the initial phage concentration.

We noted that after the 13 h time point, though the CFU counts were comparable to the uninfected controls, the *E. coli* colonies were only seen in the spots of higher dilutions (Fig. 2) while lower dilutions were clean. Notably, this effect was observed despite the viricidal treatment of the suspension with the tea extract (see Methods) to remove the extracellular phages. This result indicates that at later time points a significant fraction of PFUs was represented by infected cells, while before 11 h the free phage prevailed (and the total PFU counts were much greater).

**Fig. 2.**
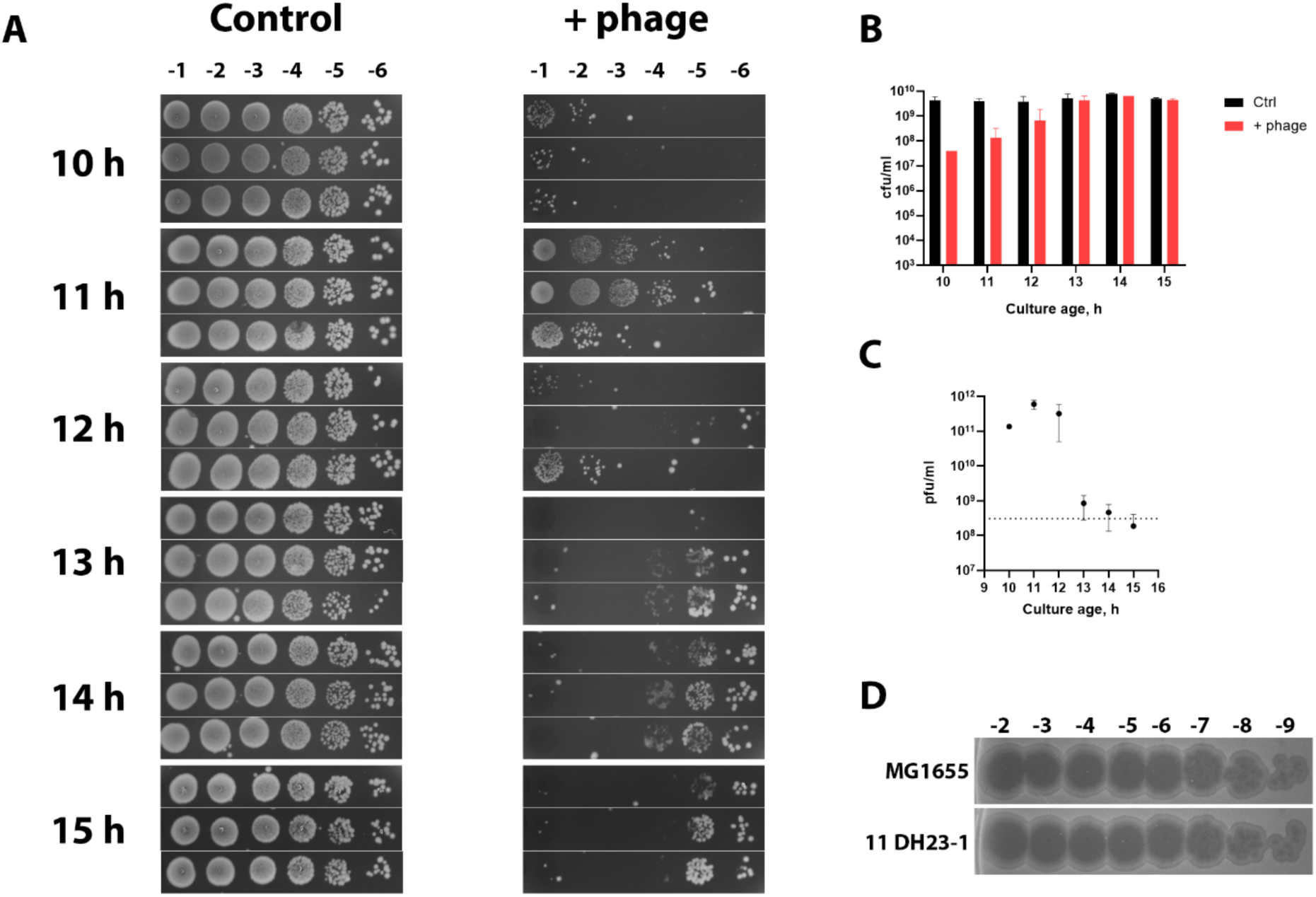
Cultures from time points 10-15h infected with DH23 phage. Cells (**A** and **B**) and phage (**C**) were titrated after 24h incubation. All time points were made in triplicate. Surviving cells from 11h culture incubated with phage can support phage propagation (**D**). The dotted line indicates the initial phage concentration at the moment of the inoculation.

We streaked out the cells grown from 11h infected cultures and tested their sensitivity to the DH23 phage. The cultures were completely sensitive showing the efficiency of plating (EOP) identical to the original strain (Fig. 2). This result is surprising because the cells surviving in the 11h infection experiment were incubated with a very high (5×10^10^ PFU/ml) phage concentration for about 24h. We conclude that a small fraction of the cells in the cultures becomes physiologically phage-tolerant but give rise to colonies with a normal phage-sensitive phenotype (see Fig. 2D); noteworthy, under the conditions tested this subpopulation does not grow that makes us to use the term “physiological phage tolerance” instead of physiological resistance. We do not know if such cells are present at earlier time points since they are masked by rising resistant population or at later time points where phage multiplication is inhibited. Further investigation of the mechanism of such phage tolerance falls outside the scope of this study.

Our results indicate that there is a clear physiological transition between early SP cells susceptible to DH23 infection and the middle SP cells which can be infected but do not support phage progeny production or release until the nutrients are added. This transition takes place within a narrow time window between 11h and 13h of culture growth (notably, the parallel cultures at 12h gave quite variable infection outcomes suggesting that this time corresponds to the transition point).

To confirm different modes of the phage DH23 interaction with the cultures before and after this transition, we compared total phage titer and infected cells (tea-resistant PFUs) in 10h and 14h infections (MOI = 0.1). The total phage titer was about 5×10^11^ PFU/ml in the 10h infection but remained at the initial level of 10^9^ PFU/ml after the infection of the 14h-old culture. At the same time the fraction of tea-resistant PFUs was less than 10% in the 10h infection and reached about 50% in the 14h infection (Fig. 3).

**Fig. 3.**
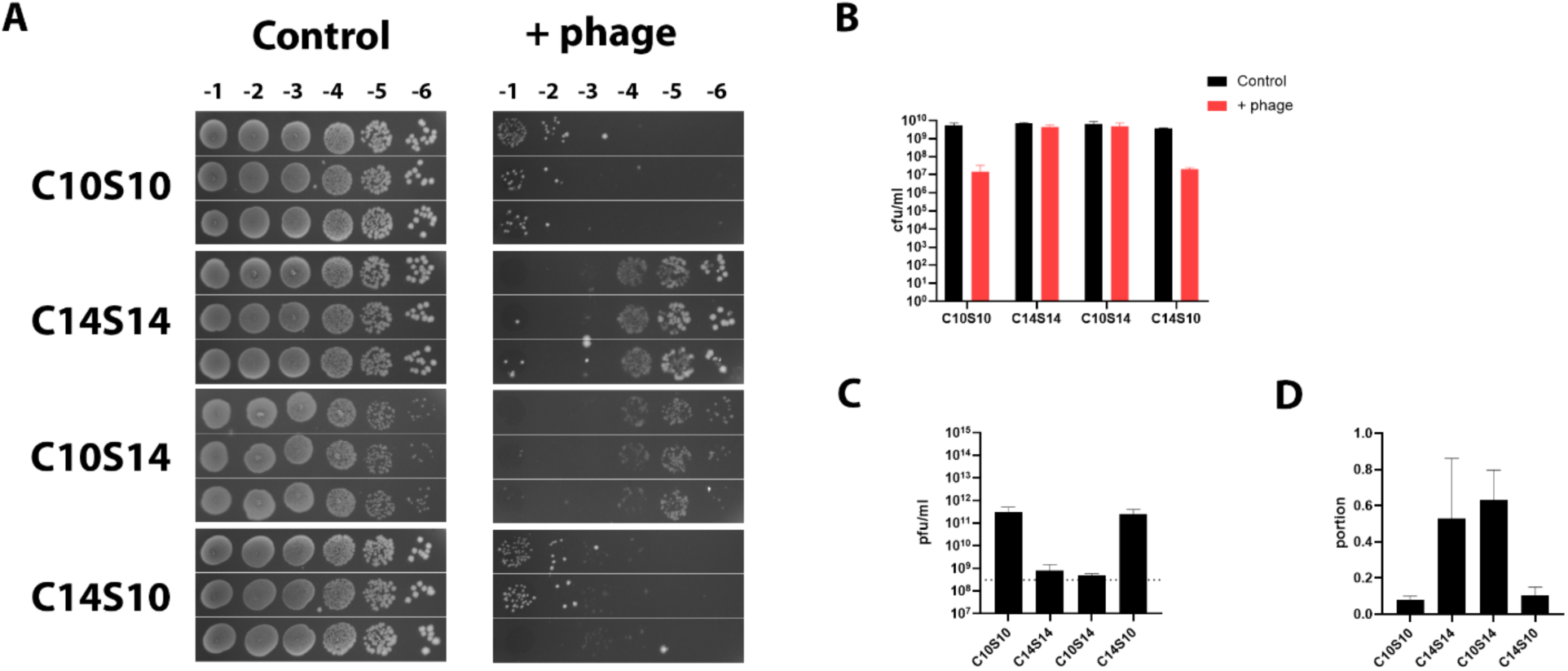
Culture medium effect on phage propagation. Cells from the 10h culture were resuspended in their own medium (C10S10) or 14h culture media (C10S14). On the other hand, cells from the 14h culture were resuspended in their own medium (C14S14) or 10h culture media (C14S10). All suspensions were incubated with phage for 24h. After incubation for 24h cells (**A**, **B**) and phage (**C**, **D**) were titrated. The dotted line indicates the initial phage concentration.

The CFU counts followed the opposite pattern: in the 14h infection they were similar to the level of the non-infected control (ca. 10^10^ CFU/ml), while at 10h viable cell counts were 4 orders of magnitude lower. Notably, as in the previous experiments, the colonies did not grow in the spots of the lower dilutions (Fig. 2) from the 14h culture but the titration looked normal in 10h infections despite much higher initial total PFU counts in these samples. These results indicate that the phage actively propagates in the 10h pre-transition culture, killing all the cells but a small fraction of tolerant ones and, potentially, an even smaller number of phage-resistant mutants. If the 14h culture is infected, the phages enter the cells but the infection stalls until these cells are transferred to fresh media on plates. This effect is similar to the outcome of T4 infection of the stationary phase cells described by E. Kutter’s group [38] though under our conditions T4 growth is completely inhibited at an earlier phase of the culture between 6h and 8h post inoculation [36]. Thus, the phage infection served as an efficient probe to reveal a previously unrecognized physiological state of the vast majority of the cells in the stationary culture which we refer to here below as middle-SP-dormancy (MSPD).

### The MSPD is induced and maintained through extracellular signaling

MSPD becomes apparent some hours after the population enters the SP, when the cells have already stopped growing and dividing. Therefore, we asked whether this effect is due to changes in the state of the medium or instead reflects the internal physiology of the cells, e.g., the stage of protein aggregate maturation recently described to influence the deepness of the bacterial dormancy [3]. To address this question, we prepared 10h and 14h host cultures, separated the cells and supernatant by low-speed centrifugation and filtered the supernatant through a 0.22 µm pore-size filter. The cultures were then re-constituted by resuspension of each of the cells in each of the supernatants, yielding four combinations. The 10h cells resuspended in 10h supernatant (the re-constituted 10h culture, referred here below as C10S10) were as sensitive to the phage DH23 as the initial 10h culture. The same was true for the C14S10 variant. At the same time both C10S14 and C14S14 showed blocked infection (Figs 2, 3). We concluded that 14h supernatant (spent media) was responsible for MSPD induction and maintenance. The effect of the supernatant was also easily detectable in the experiments monitoring the infected culture OD600 (Fig. 4), which can be used instead of PFU and CFU titrations to test the influence of different conditions on MSPD.

**Fig. 4.**
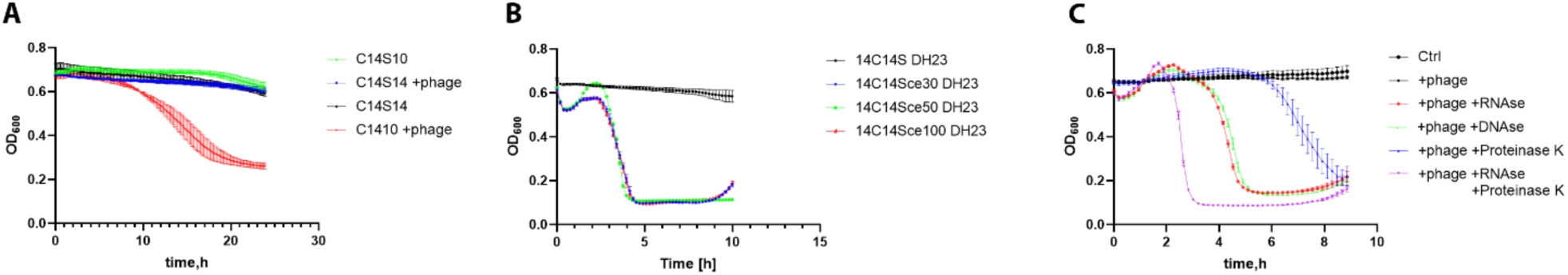
Influence of the culture spent media modifications on the phage DH23 infection of the SP culture. (**A**) Cells from 14h culture were resuspended in their own spent medium (C14S14) or 10h culture media (C14S10). (**B**) Cells from 14h culture were resuspended in their own medium filtered through centrifuge concentrators with cutoffs of 30 (C14S14ce30), 50 (C14S14ce50) and 100 kDa (C14S14ce100). (**C**) Enzymes were added to the 14h culture.

We then asked whether the effect of the 14h supernatant was due to the lack of some factor essential for phage multiplication or can be explained by the presence of an inhibitor. We repeated the experiment described above using different treatments of the 14h supernatant which may potentially remove or destroy the signal. We filtered S14 through centrifuge concentrators with cutoffs of 30, 50 and 100 kDa, added RNAse, DNAse or proteinase K. In the case of the proteinase K a separate test was performed to ensure that phage infectivity is not compromised by the protease treatment under the conditions of our experiment (data not shown). Surprisingly, all the treatments applied removed the inhibitory activity of S14 (Fig. 4) allowing phage multiplication and efficient lysis of the bacterial culture.

The RNAse and DNAse showed similar efficacy; the effect of proteinase K was weaker and appeared additive to that of the RNAse. Filtration through the 100 kDa cutoff membrane (as well as 30 and 50 kDa cutoffs) also efficiently removed the MSPD-maintenance signal.

We then repeated the experiment with infection of the 14 h culture with the phage DH23 with and without RNAse treatment. The bacteriophage production was highly elevated in the RNAse-treated culture and all the PFUs were represented by free phage particles (tea-sensitive). In the control 14h culture (without the RNAse treatement) more than half of the PFUs (Fig. 5) were due to infected cells (tea-resistant). Interestingly, in both treated and untreated cultures CFU counts were similar to the uninfected (phage-free) control but the titration of the untreated infected culture (tea plus phages) produced the above-mentioned effect of the colony growth inhibition in lower dilutions due to lysis by phages released from the infected cells. In the RNAse-treated culture, the high cell density was due to secondary re-growth of phage-resistant cells and therefore the CFU counts looked normal (Fig. 5).

**Fig. 5.**
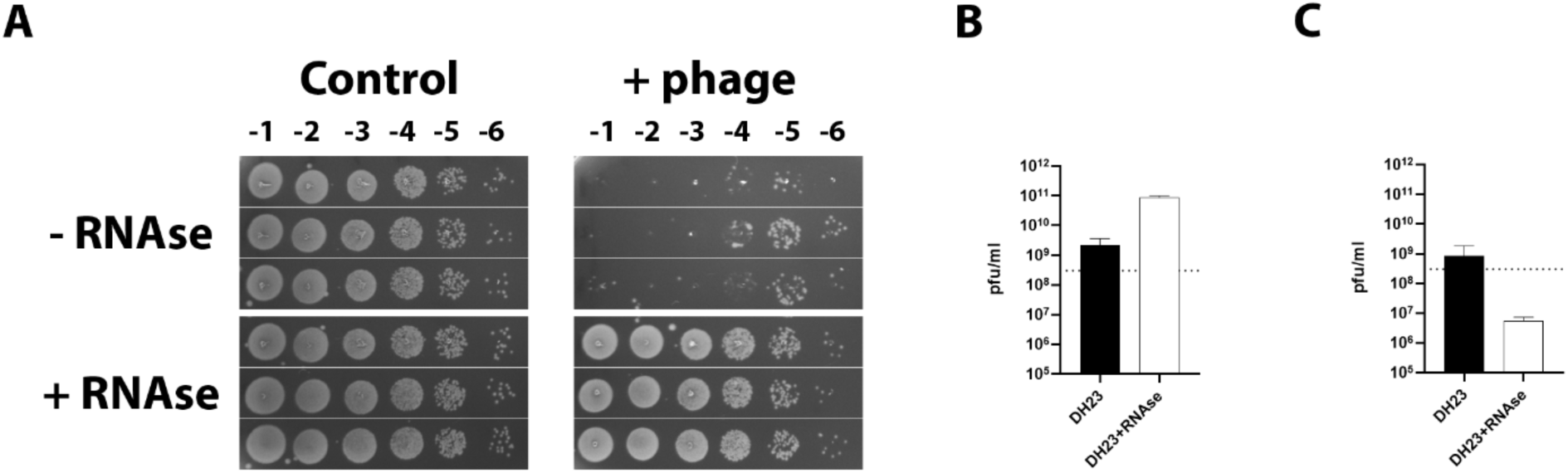
Phage DH23 multiplication and cell-killing activity in RNAse-treated 14h culture. (**A**) CFU counts (triplicated results shown), (**B**) Total PFU counts, and (**C**) tea-resistant PFU counts. All the titrations were performed after 24h incubation with phage. Dotted lines in panels B and C indicate the initial phage concentration.

We concluded that MSPD is induced by extracellular particles smaller than 0.22 µm but larger than 100 kDa filter pores (approximately 10 nm). This complex includes both DNA and RNA which are necessary for its activity. Most probably it also contains protein(s) which may protect nucleic acids from naturally present nucleases. Here below we term this putative complex the middle-SP signal particle (MSP). The removal or enzymatic destruction of MSP makes the cells suitable for the productive DH23 infection.

### The effect of MSP on infection of other SPI-bacteriophages

Using OD600 monitoring in the microplate reader-incubator, we assessed the RNAse treatment effect on the infection of the 14h *E. coli* MG1655 culture by three other SPI phages described by us previously [36] including phage T7 and our own isolates. All the phages caused efficient lysis of the 14h culture if the RNAse was added. No OD600 drop was observed without the RNAse treatment, indicating that MSPD blocked all the phages. The RNAse treatment had little or no effect on the culture OD in the absence of phage infection (Fig. 6).

**Fig. 6.**
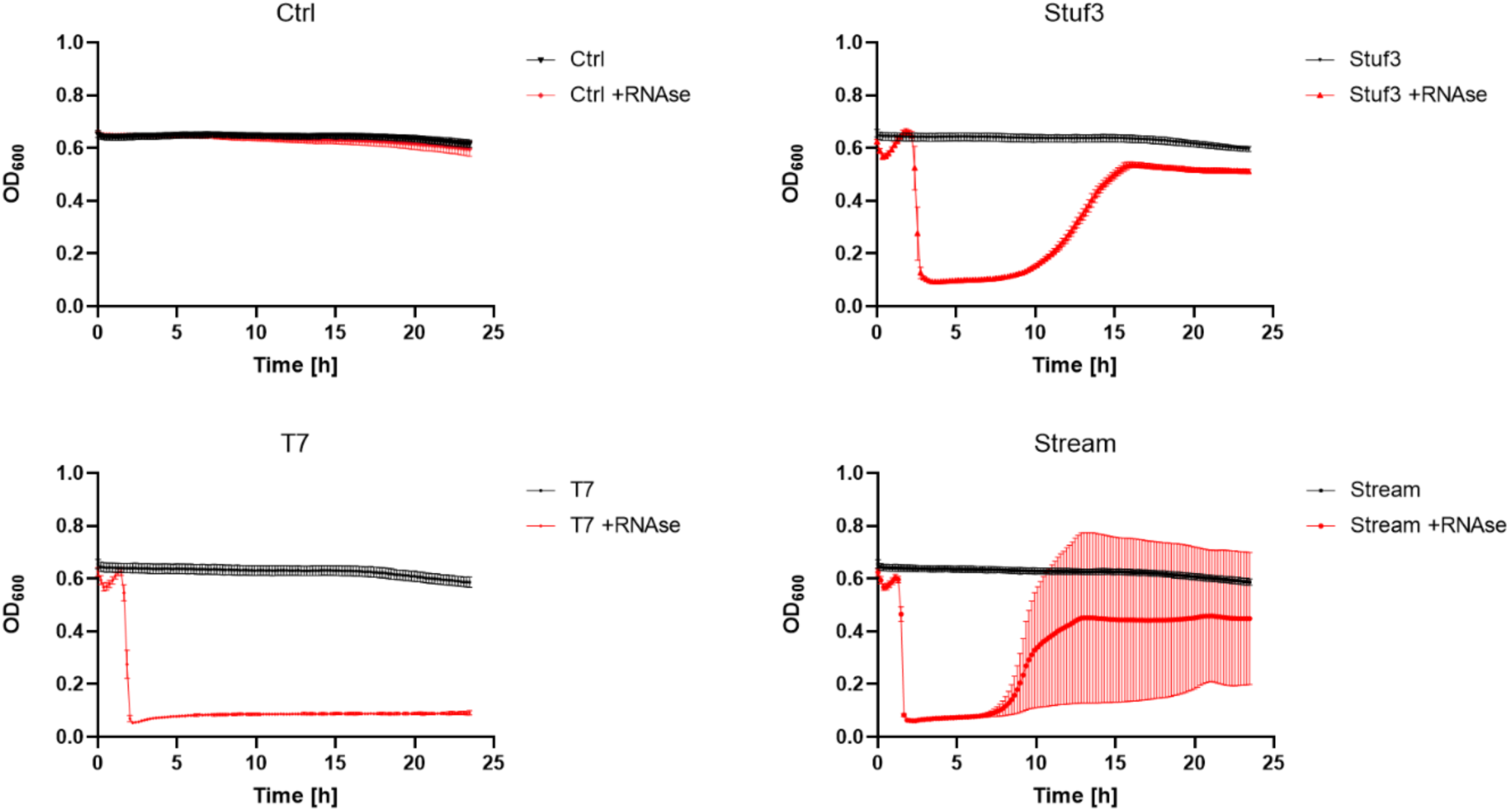
Lysis of the 14h culture by SPI phages with or without RNAse treatment.

### MSPD lifting influences cell morphology and antibiotic sensitivity

In order to test the effect of the MSP signal removal on cell physiology, we first compared the morphology of the cells in the 14h culture treated with RNAse for 1.5 h with the control culture to which an equal amount of RNAse buffer was added. The cells in the treated culture were found to have increased in both length and diameter compared to the non-treated cells (Fig. 7). This reversal of the morphology changes was not associated with any significant increase in DNA synthesis as determined by Click-iT staining (Fig. 7). Also, we did not observe any increase in cell number after RNAse treatment (Fig. 7).

**Fig. 7.**
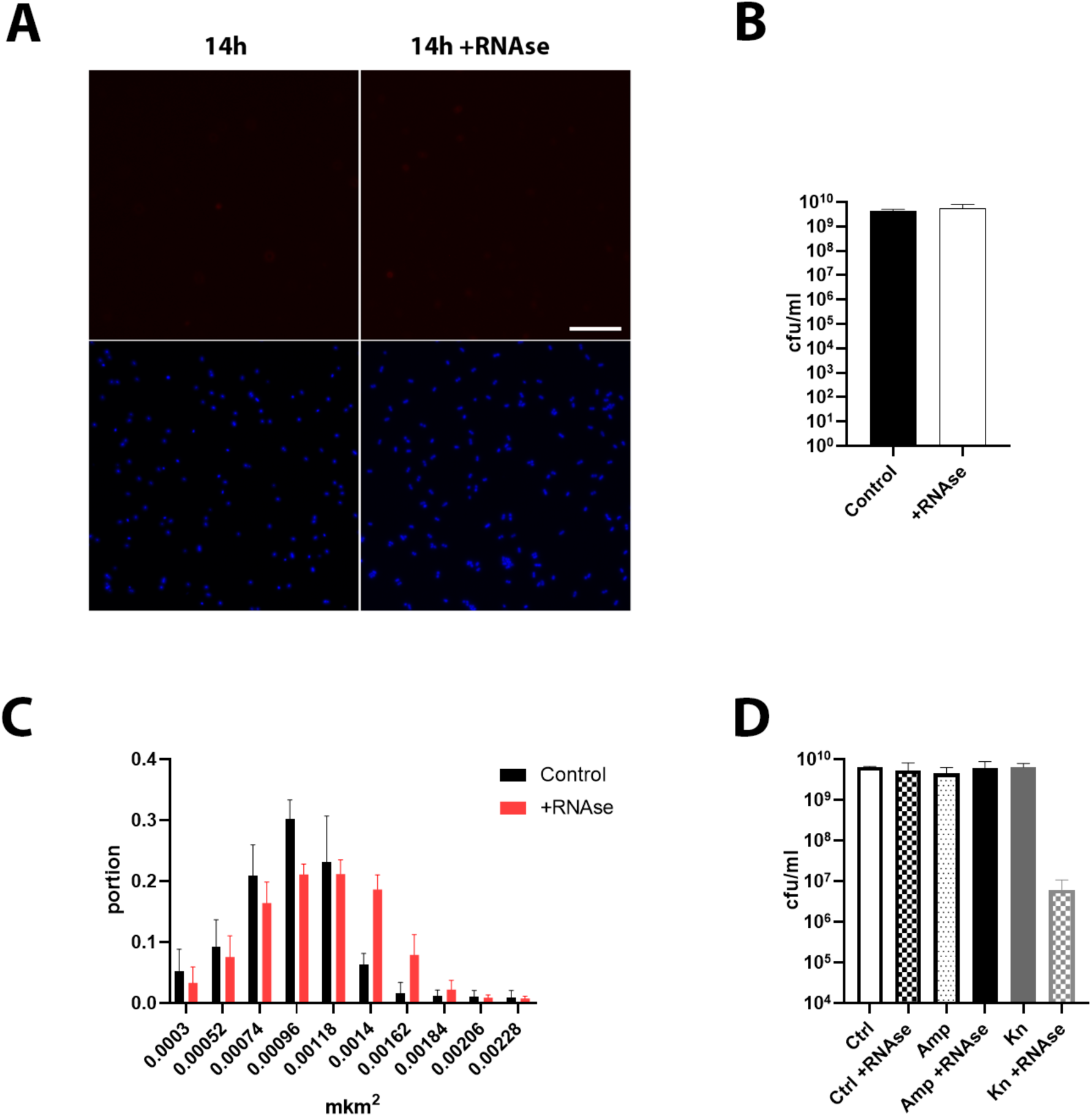
Reactions of the *E. coli* cells in 14 h culture to the RNAse treatment. (**A**) 14h culture incubated with or without RNAse for 1.5 h more and stained with Click-iT EdU. (**B**) RNAse treated culture was titrated. (**C**) Size (projection area) distribution of the cells in RNAse-treated and non-treated 14h culture. (**D**) Killing effect of ampicillin and kanamycin on 14h culture with or without the RNAse treatment.

We speculate that enlargement of the cells would require peptidoglycan remodeling and protein synthesis. These processes may increase the sensitivity of the culture to the antibiotics blocking the corresponding processes such as ampicillin, a peptidoglycan synthesis inhibitor, or kanamycin, distorting protein synthesis and causing cell damage by toxic aberrant peptides.

We compared the CFU counts in 14h culture treated with each of the antibiotics for 4h with or without the RNAse treatment. As expected, the intact SP culture was tolerant to both ampicillin and kanamycin. The RNAse treatment did not affect the ampicillin sensitivity but caused a marked enhancement of the kanamycin-induced killing, with CFU counts dropping 3 orders of magnitude compared to the control (Fig. 7). These results indicate that the activation of the peptidoglycan remodeling, even if present, was not sufficient to support ampicillin-induced cell wall damage to the level needed for the cell lysis. At the same time MSP removal and MSPD lifting activate protein synthesis making the cells sensitive to kanamycin even without active growth and multiplication.

### MSP role in SP establishment

We noted that the re-growth of resistant mutants was inhibited when the DH23-mediated lysis of the early SP cultures occurs before the MSPD onset (5 – 11h or the reconstituted C14S10 culture; Figs. 2 and 3). However, the re-growth is enabled in the RNAse-treated or DNAse-treated 14 h cultures (Figs. 4–6). These results suggest that the MSP accumulation begins earlier during the culture growth prior to the MSPD. To investigate how MSP-mediated effects influence the growth of the *E. coli* culture we followed the growth curves of the cultures with or without RNAse added simultaneously with the culture inoculation. In the absence of RNAse, the growth rate started to slightly decline from 3.5 h (Fig. 8) and the culture reached the SP by 5 h (in the conditions of growth in 96-well plate in the reader-incubator device). The culture treated with any of the nucleases continued to grow at the same rate for an additional hour, reaching a ca. 30% higher density. Interestingly, in contrast to the normal 14h culture treated with RNAse, the morphology of the cells after 16h of growth in the presence of RNAse was not distinguishable from the control (data not shown). These results indicate that MSP signaling may allow the bacterial population to switch cooperatively into the resource-saving mode proactively before the medium is fully exhausted.

**Fig. 8.**
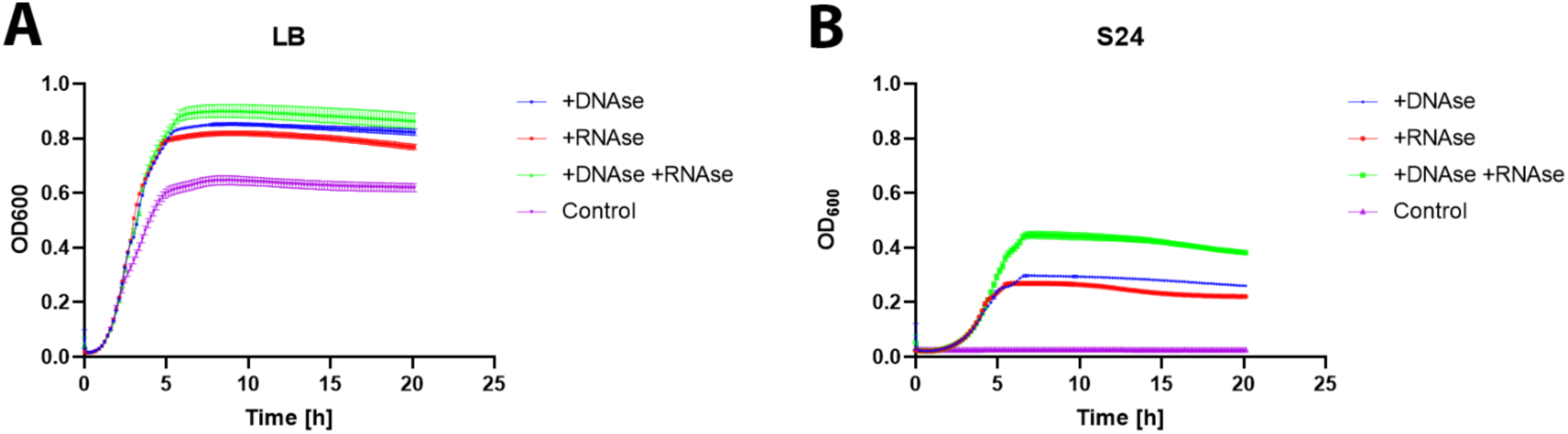
Growth of *E. coli* MG1655 in fresh LB or spent medium from 24h culture with or without nuclease supplementation. Cells from 24h culture were resuspended in 0.9% NaCl, 500-fold diluted LB or 24h culture medium, with or without nucleases.

To test this hypothesis, we prepared spent medium (supernatant) from a 24 h culture that had reached late SP (as defined in [20]) and inoculated it with the 24 h culture cells resuspended in physiological saline. The same cultures were set also in fresh LB media. These experiments were performed with and without RNAse and DNAse added simultaneously with the inoculation. In the absence of any externally added nuclease, the cells are not able to resume growth in 24 h spent media. However, if the MSP activity is destroyed by the nucleases, the growth is initiated with a slightly longer lag-phase and the culture reaches the OD600 of about 60% of the SP culture grown in the fresh LB media. In fresh LB the addition of DNAse or RNAse increased the OD600 plateau level about 1.5 times (Fig. 8). These results indicate that cell division in conventional cultures is stopped in a coordinated manner prior to medium exhaustion.

## Discussion

Testing of activity, such as the ability to form plaques, EOP or efficiency of phage adsorption and/or multiplication on cultures of different host strains under different conditions, is extensively used to discover and analyze particular mechanisms of bacteriophage life cycle or phage-host interactions. This research formed the grounds for deciphering the biological nature of viruses and revealing basic molecular mechanisms of life (recently reviewed in [39]). Phages may also be employed as probes to reveal specific properties of host cells. This includes high-resolution strain differentiation with phage typing [40–42], studying the architecture of the cell surface [43,44], probing the protective efficiency of external structures such as the O-antigen [45–47], and identification of bacterial phase variations [48]. Here we used testing of the interactions of SPI-coliphages with the standard laboratory *E. coli* strain MG1655 as a tool which helped to reveal a fine periodization of the physiological state of *E. coli* within the previously defined stationary phase of growth.

In most of the works the regulation of the general stress response and other physiological switches associated with the *E. coli* culture entry into the SP (see Introduction section and refs cited there) are described as a cellular-level response to starvation, intoxication by metabolic products and other stressors. Here we demonstrated that both population growth arrest and the development of the previously unrecognized MSPD state are coordinated through intercellular communication mediated by an unusual extracellular MSP signal.

Our results indicate that MSP-mediated growth arrest saves about 30-40% of the initial medium resources from being wasted on a further increase in the density of the population, which would then run out of the energy supply necessary for the maintenance of the viability of the cells [49] over an extended period of time, enabling the population to survive until nutrients become available again. Interestingly, the MSP-dependent pre-emptive growth arrest and the MSPD state, effectively sensed by the phage DH23 and, probably, by some other SPI phages (see [36]), are two distinct physiological switches. In our experiments RNAse or DNAse treatment of the 14 h culture lifted MSPD and restored bacteriophage and kanamycin sensitivity (Fig. 4–7), but these treatments did not enable further population growth. At the same time the inoculation of the RNAse-treated 24h spent medium with a small number of cells resulted in considerable bacterial growth (Fig. 8).

These observations indicate that there is another population-density dependent mechanism controlling *E. coli* proliferation which is in agreement with the observations by Ughy et al. [50] who defined a minimal stationary cell concentration (MSCC) point, down to which the SP bacterial cultures do not resume growth upon dilution. Notably, in our experiments bacterial cultures grown in LB supplemented with RNAse reached a much higher density compared to the control suggesting that the second hypothetical mechanism is mainly responsible for preventing bacterial regrowth in too dense populations but not for stopping the actual growth at some defined level, which the MSP signaling appears to be responsible for. Notably, in the phage-lysed early SP cultures (Fig. 1-2) the secondary growth of the resistant mutants is hindered by the remaining MSP, however, in the SP cultures treated with RNAse and then lysed with the phage the resistant mutants readily re-establish a high-density population indicating that the either the MSCC effect is controlled by a different signal that can is removed during the phage-mediated lysis (but not by the RNAase treatment, see the Ctrl panel at the Fig. 6) or the living cells themselves are essential for the MSCC sensing.

From the eco-evolutionary perspective, the MSCC-MSP mechanism may represent an effective strategy for collective hedging of the risk to run out of the nutrients necessary for the viability maintenance in SP. In such a case this strategy it may be compromised if a certain number of cheater cells, mutant or physiologically non-responsive to MSP signaling, emerge in the population. However, in our experiment we did not observe such an effect (e.g. no cases of the spontaneous culture overgrowth to the plateau level seen in the RNAse supplemented media, Fig. 8) though our current study was not focused on the ecology and microevolution of the SP populations).

The unusual nature of the MSP, which appears to be a large complex including both RNA and DNA required for its activity, creates diverse possibilities for modulation of the SP population physiological state by nucleases or other MSP-inactivating enzymes released from dying cells or provided externally (e.g. by a macro-host of the symbiotic microbial population). The enzymes altering the physiological state of the neighboring cells may also be released or produced due to the phage lysis of a fraction of this population or of some co-habiting cells of different microbial strains or species, thus creating a window for phage-driven alterations of the physiological state of the non-infected cells (earlier named by us as POSE effects, [51], see also [52]). We also demonstrated that the artificial lifting of MSPD leads to an increase in sensitivity to some antibiotics to which the SP population is tolerant. This effect may be investigated to develop improved clinical protocols to harness potential nuclease-antibiotic synergy against the SP populations for more efficient treatment of chronic infections which often require long antibiotic courses despite the *in vitro* sensitivity of the causative agent to the drug.

For the moment we are not able to provide any mechanistic insights into MSP formation and mode of action on the cell. However, the currently available data suggest that these mechanisms are fundamentally different from known QS systems and other types of intercellular communication in bacteria.

## Materials and methods

### Bacterial and phage strains and their cultivation

The standard *E. coli* K-12 MG1655 strain and bacteriophage T7 were from our laboratory collection. Other bacteriophages were isolated by us previously from various environmental objects. For details see [36,37]. Bacterial cultures were grown on LB media (10 g Bacto-Tryptone, 5 g yeast extract, 10 g NaCl, distilled water up to 1 L), supplemented with 15 g of Bacto-agar per 1 L for the plates or 6 g/L for the soft agar. Bacteriophage titration was performed using standard double-layer plating. For bacteriophage cultivation, mid-log phase *E. coli* MG1655 culture with OD600 of 0.4 – 0.5 was inoculated with the phage at MOI 0.01 and incubated at 37°C with agitation at 250 rpm until visible lysis (typically 3–4 h). Then an 0.02 vol of chloroform was added and the lysate was cleared by centrifugation at 12 000 g for 5 min.

For preparation of standardized *E. coli* cultures of different age (post-inoculation time) fresh overnight culture was prepared and kept at 4°C during the working day as inoculum; the culture was briefly vortexed before use. For each time point a separate 200 ml Erlenmeyer flask with 50 ml of LB media was prepared and cooled down in the fridge. To start the culture 100 µl of the cold inoculum was added per flask (1:500 v/v) and the flasks were incubated at 37°C with agitation at 250 rpm for the defined number of hours indicated in the assays. Importantly, each flask was opened only once; independent cultures were used for each time point in triplicate.

In some experiments the cells and supernatant of the cultures were separated by centrifugation at 7000 g for 15 s and the supernatant was filtered through a 0.2 µm pore size syringe filter. Then the cultures were re-constituted by resuspension of the cells in the same volume of the supernatant from the same or another culture (see the Results section). Also, some tests were performed with the supernatants filtered through centrifuge membrane concentrators (Sartorius, UK) with a 100 kDa cutoff.

### Enzymes and antibiotics

In some experiments the cultures or media were supplemented with 2 µg/ml of RNAse A (Biolabmix, Russia) or 1 U/ml of DNAse I (Thermo Fisher Scientific, USA) or 4 µg/ml of proteinase K (ServiceBio, China) or combinations of these enzymes. Ampicillin and kanamycin treatments were used in standard concentrations (100 µg/ml and 50 µg/ml, respectively).

### Viricidal tea extract preparation

The viricidal extract was prepared as described in [53]. Briefly, 10 g of black leaf tea (“The crown of Russian Empire”, Russia) were poured with 100 ml of boiling deionized water and then incubated at 55°C for 30 min. The extract was filtered through a sterile filter paper. Tea extract was kept at 4°C and remained active for 1 month. A precipitate that forms after the cooling down of the extract was dissolved by heating before each use.

### Bacteriophage replication test

To test phage replication in the cultures of different ages 5 ml of the culture was inoculated with the phage DH23 at a multiplicity of infection (MOI) of about 0.01 PFU/CFU. The cultures were then cultivated at 37°C with agitation at 250 rpm for 24 h. The uninfected controls were always set up. After the cultivation, serial dilutions of the cultures were spotted on the host lawn to enumerate the total phage titer. For CFU titration the first dilution was made in viricidal tea extract, incubated for 10 min at room temperature and then the subsequent dilutions were prepared in the LB media. For the infected-cell PFU titration the same treatment was used with dilutions plated on bacterial lawn.

### Click-iT

100 µl of cell culture was incubated with 1 mM EdU for 15 min at 37°C and washed twice with PBS buffer. Washed cells were fixed by adding 100 µl of 6% paraformaldehyde solution in PBS. Fixed cells were stained according to the manufacturer’s instructions.

### Microscopy

Axiovision M1 microscope (Zeiss, Germany) equipped with a 100x lens and an Axiocam 503 mono Zeiss camera was used to capture images. Images were analyzed with ImageJ software.

### Growth curves

Cell growth and degradation curves were recorded in 96-well format at 37°C with agitation at 250 rpm using an Infinite M Plex plate reader (Tecan instruments, USA). *E. coli* cultures were loaded at 100 µl per well. Each experimental point was made in triplicate. OD600 measurements were collected for 15-20 h at 10-minute intervals.

For the determination of the growth curves in LB and 24 h culture spent media with or without nucleases, the cells from the 24 h culture were collected by centrifugation at 7000 g for 15 s and resuspended in the initial volume of the physiological saline (0.9% NaCl). This suspension was used to inoculate 1:100 (v:v) fresh LB media or the spent media (filtered supernatant) from the 24 h culture, with or without RNAse and/or DNAse added. The curves were then recorded as described above.

## Supporting information

Figure S1. Growth curve of E. coli MG1655 strain

## Acknowledgements

The authors are grateful to Stephen Abedon from Ohaio State University, USA, for the critical reading and linguistic editing of the manuscript))

