## Supplementary figures and images for "Bacteriophage infection reveals pre-emptive cooperative dormancy in stationary phase *Escherichia coli*"

### Figure S1. Growth curve of E. coli MG1655 strain

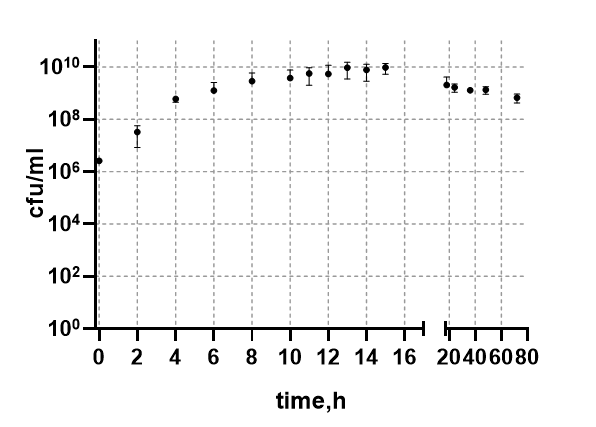
